# An antioxidant screen identifies ascorbic acid for prevention of light-induced mitotic prolongation in live cell imaging

**DOI:** 10.1101/2022.06.20.496814

**Authors:** Tomoki Harada, Shoji Hata, Masamitsu Fukuyama, Takumi Chinen, Daiju Kitagawa

## Abstract

Phototoxicity is an important issue in fluorescence live imaging of light-sensitive cellular processes such as mitosis, especially in high spatiotemporal resolution microscopy that often requires high-intensity illumination. Among several approaches to reduce phototoxicity, the addition of antioxidants to the imaging media has been used as a simple and effective method. However, it remains unknown what are the optimal antioxidants that could prevent phototoxicity-induced defects during mitosis in fluorescence live cell microscopy. In this study, we analyzed the impact of phototoxicity on the mitotic progression in fluorescence live imaging of human diploid cells and performed a screen to identify the most efficient antioxidative agents that reduce it. Quantitative analysis shows that high amounts of light illumination cause various mitotic defects such as prolonged mitosis and delays of chromosome alignment and centrosome separation. Among several antioxidants known to reduce cellular phototoxicity, our screen reveals that ascorbic acid significantly alleviates these phototoxic effects in mitosis. Furthermore, we demonstrate that the addition of ascorbic acid to the imaging media enables fluorescence imaging of mitotic events at very high temporal resolution without obvious photodamage. Thus, this study provides a simple and practical method to effectively reduce the phototoxic effects on mitotic processes in fluorescence live cell imaging.

## Introduction

Live cell imaging is a powerful technique for studying dynamic cellular processes. Recent advances in fluorescence imaging have enabled us to visualize molecular dynamics with high spatiotemporal resolution (Sung et al., 2011). However, the application of such high-resolution imaging on living cells is limited due to phototoxicity caused by high-intensity or prolonged sample illumination. Phototoxicity occurs through the light-induced formation of reactive oxygen species (ROS) (Laissue et al., 2017; Icha et al., 2017), which react with numerous cellular components and disrupt their structures and functions. Severe oxidization results in cellular abnormalities such as membrane blebbing and subsequent cell death (Zdolsek et al., 1990; Tosheva et al., 2020). Even without apparent changes in cellular morphology, high amounts of light illumination can cause substantial cell dysfunction such as abnormal intracellular calcium homeostasis (Roehlecke et al., 2009; Nishigaki et al., 2006). Therefore, reducing phototoxicity in fluorescence live cell imaging is an important factor to accurately capture cellular processes with high spatiotemporal resolution and to avoid misinterpretation of experimental results.

Mitosis is a series of dynamic processes through which a parent cell divides into two genetically identical daughter cells (Walczak et al., 2010). Live imaging of these mitotic events however is not trivial, as they are known to be particularly sensitive to phototoxicity. Excitation light, especially at a wavelength of 488 nm, has been shown to inhibit mitotic entry in a dose-dependent manner (Douthwright and Sluder, 2017), and to cause prolongation of mitotic duration (Schilling et al., 2012). Since cells round up and increase their height during mitosis (Taubenberger et al., 2020), three-dimensional (3D) imaging that covers the entire cell over a wide z-axis range is necessary to capture the spatial dynamics of all relevant cellular components in this process. Furthermore, high temporal resolution is required to record the highly dynamic and complex nature of mitosis, which is completed in less than an hour. However, 3D time-lapse imaging with short intervals requires a very high frequency of excitation light exposure, which can induce phototoxicity in living cells.

Several approaches have been proposed to decrease photodamage in fluorescence imaging of living cells, including the use of advanced microscopes with low phototoxicity such as light-sheet microscopes (Tosheva et al., 2020). However, these microscopes are often expensive and difficult to implement. The use of excitation light at a longer wavelength can also reduce cellular phototoxicity, although the effective brightness of fluorescent proteins excited at longer wavelengths, such as near-infrared fluorescent proteins, is much lower than that of green or red fluorescent proteins in mammalian cells (Icha et al., 2017). Adding antioxidants to the live cell imaging buffer is another approach to reduce cellular phototoxicity by scavenging ROS and limiting oxidative stress upon light illumination (Icha et al., 2017). The antioxidants sodium pyruvate and Trolox have been shown to protect cells from light-induced cell death (Natoli et al., 2016) and G2 arrest, respectively (Douthwright and Sluder, 2017). Although the use of antioxidants in the live cell imaging buffer is a simple and effective approach to generally reduce photodamage, the optimal antioxidative agents for the specific aim to accurately capture mitotic dynamics in fluorescence microscopy remain unclear.

In this study, we performed a screen to identify antioxidants that are capable of alleviating light-induced mitotic abnormalities in fluorescence live imaging. Among several compounds, ascorbic acid, also known as vitamin C, significantly reduces photodamage in mitotic cells without showing indications of cytotoxic side-effects. We demonstrate that at the appropriate concentrations required to reduce mitotic phototoxicity, ascorbic acid does not perturb cell survival or the accurate segregation of the chromosomes. Moreover, the addition of ascorbic acid to the imaging media enables time-lapse 3D imaging of mitotic processes at very short temporal intervals, which has been so far difficult to achieve without obvious photodamage. Therefore, this study provides an effective solution for reducing the mitotic phototoxicity and demonstrates its application for fluorescence imaging of mitotic dynamics at very high spatiotemporal resolution in living cells.

## Results

### High light illumination causes abnormal prolongation of mitosis and delays chromosome alignment and centrosome separation in live cell 3D imaging

We first investigated the vulnerability of the mitotic processes to phototoxicity in fluorescence live cell imaging of RPE1 cells, which are one of the most widely used non-transformed human diploid cells. To this end, we used RPE1 cells stably expressing mNeonGreen (mNG)-fused Histone H2B and mRuby2-fused γ-tubulin to visualize the dynamics of chromosomes and centrosomes, respectively (Hata et al., 2019), which allowed us to observe key events in the mitotic progression. To analyze photodamage in mitosis, we set two different conditions of excitation laser light illumination referred to as low and high conditions (Fig. 1a). In the high condition, we used 488 nm wavelength excitation light with >6 times higher laser power and 4 times longer exposure time compared to the low condition. The setting for the 561 nm wavelength excitation light was the same in both. Under each condition, z-stack images (21 planes with 1 μm z-steps) of the RPE1 cells were taken using a spinning disk confocal microscope in time-lapse at 3-minute intervals for a total duration of 12 hr.

**Fig.1:**
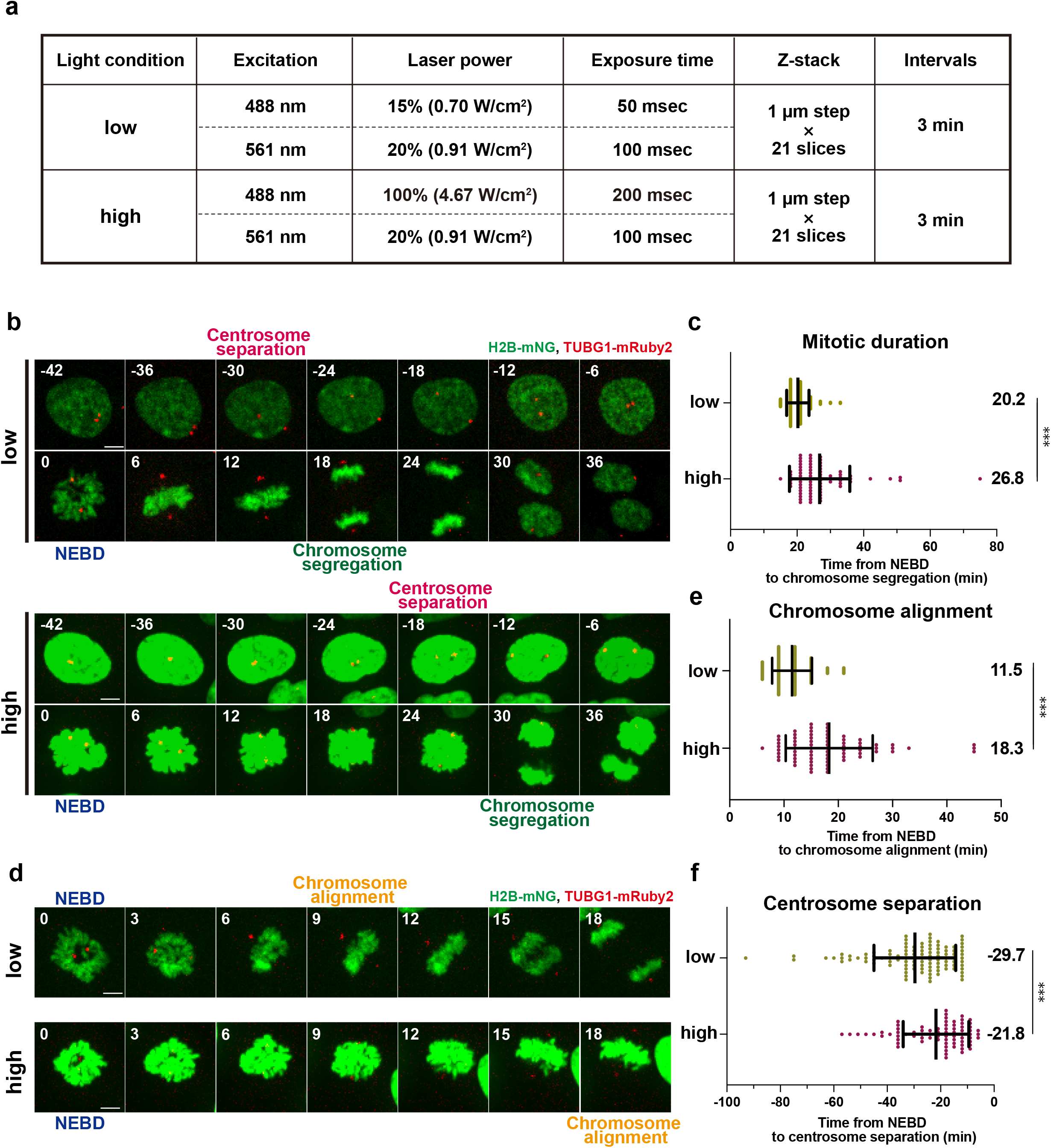
Intensive light exposure induces mitotic prolongation and delays chromosome alignment and centrosome separation in live cell imaging. **a,** The imaging conditions used for fluorescence live cell imaging. Apart from the variable intensity and exposure time of the 488-nm excitation light, all other settings are the same between the low and high conditions. **b,** Time-lapse imaging of mitotic RPE1 cells stably expressing H2B-mNG and TUBG1-mRuby2 in the low and high conditions (3-min imaging intervals). Representative still images with different settings of brightness and contrast are shown due to the intense illumination in the high condition. T=0 is designated as the time point of nuclear envelope breakdown (NEBD, time shown in min). The timing of centrosome separation and chromosome segregation are indicated. Scale bar, 5 μm. **c**, Quantification of mitotic duration from **b**. The time from NEBD to chromosome segregation was measured. n>70 cells from three independent experiments. **d**, Time-lapse images from NEBD to chromosome alignment in the low and high conditions (3-min intervals) from the same samples depicted in **b**. Representative still images are shown as in **b**. **e**, Quantification of the time required for chromosome alignment from **d**. The time from NEBD to chromosome alignment at the metaphase plate was measured. n>70 cells from three independent experiments. **f**, Quantification of the timing of centrosome separation from **b**. The time from centrosome separation (defined by >4 μm inter-centrosomal distance) to NEBD was measured. n>70 cells from four independent experiments. In **c**, **e**, and **f**, data are mean ± S.D., and *P* values were calculated by Mann-Whitney *U*-test. ***P < 0.001.

Using this setup we observed that the timing of chromosome segregation was delayed in the mitotic cells exposed to high illumination (Fig. 1b). To compare mitotic duration, the time from nuclear envelope breakdown (NEBD) to chromosome segregation was measured for each condition. As in previous reports, the duration of mitosis in RPE1 cells was approximately 20 min in the low light condition (Yang et al., 2008; Hata et al., 2019) (Fig. 1c). In contrast, cells exposed to the high-light illumination took a significantly longer time to initiate chromosome segregation after NEBD. This data is consistent with the previous evidence that strong illumination of blue light prolongs the duration of mitosis in RPE1 cells (Schilling et al., 2012). Further analysis of chromosome dynamics revealed that the time from NEBD to chromosome alignment at the metaphase plate was prolonged in the high condition (Fig. 1d, e), indicating that chromosome congression is particularly sensitive to illumination with blue light. We also found that centrosome behavior before NEBD was affected in the high condition (Fig. 1b). At the onset of mitosis, the two centrosomes are separated from each other at precise timing to initiate the formation of a bipolar spindle (Kaseda et al., 2012; Hata et al., 2019). When NEBD was used as a reference for mitotic events, centrosomes were separated at 29.7 min and 21.8 min on average before NEBD in the low and high conditions, respectively (Fig. 1b,f). This data shows that the timing of centrosome separation is delayed by high-light illumination. It should be noted that, even under the high condition, neither cell death nor other severe abnormalities were observed, suggesting that these mitotic events are specifically sensitive to phototoxicity. Taken together, these results indicate that high amounts of blue light illumination in live cell imaging cause various mitotic abnormalities.

### Shortening the total acquisition time by cell-cycle synchronization does not alleviate the mitotic phototoxicity in live cell imaging

During the live cell imaging, cells entered mitosis at various time points due to the asynchronous culturing condition. Given that the accumulation of ROS causes cellular abnormalities (Hoebe et al., 2007), cells that enter mitosis at later time points during the fluorescence imaging are expected to exhibit stronger mitotic phototoxicity than those that initiate division at earlier time points. In fact, a positive correlation between the duration of mitosis and the timing of mitotic entry after the start of live imaging was observed under the high-light illumination (Fig. 2a, b). In particular, the cells that entered mitosis in the first half of the acquisition (within 6 hr from the onset of imaging) showed a shorter duration of mitosis compared to those in the latter half (6-12 hr), and even those in the total imaging time (0-12 hr) (Fig. 2c). This data suggests that the extent to which cells are exposed to blue light before entering mitosis is an important factor in the occurrence of the mitotic phototoxicity and that reducing the pre-mitotic light exposure time would alleviate the phototoxicity.

**Fig.2:**
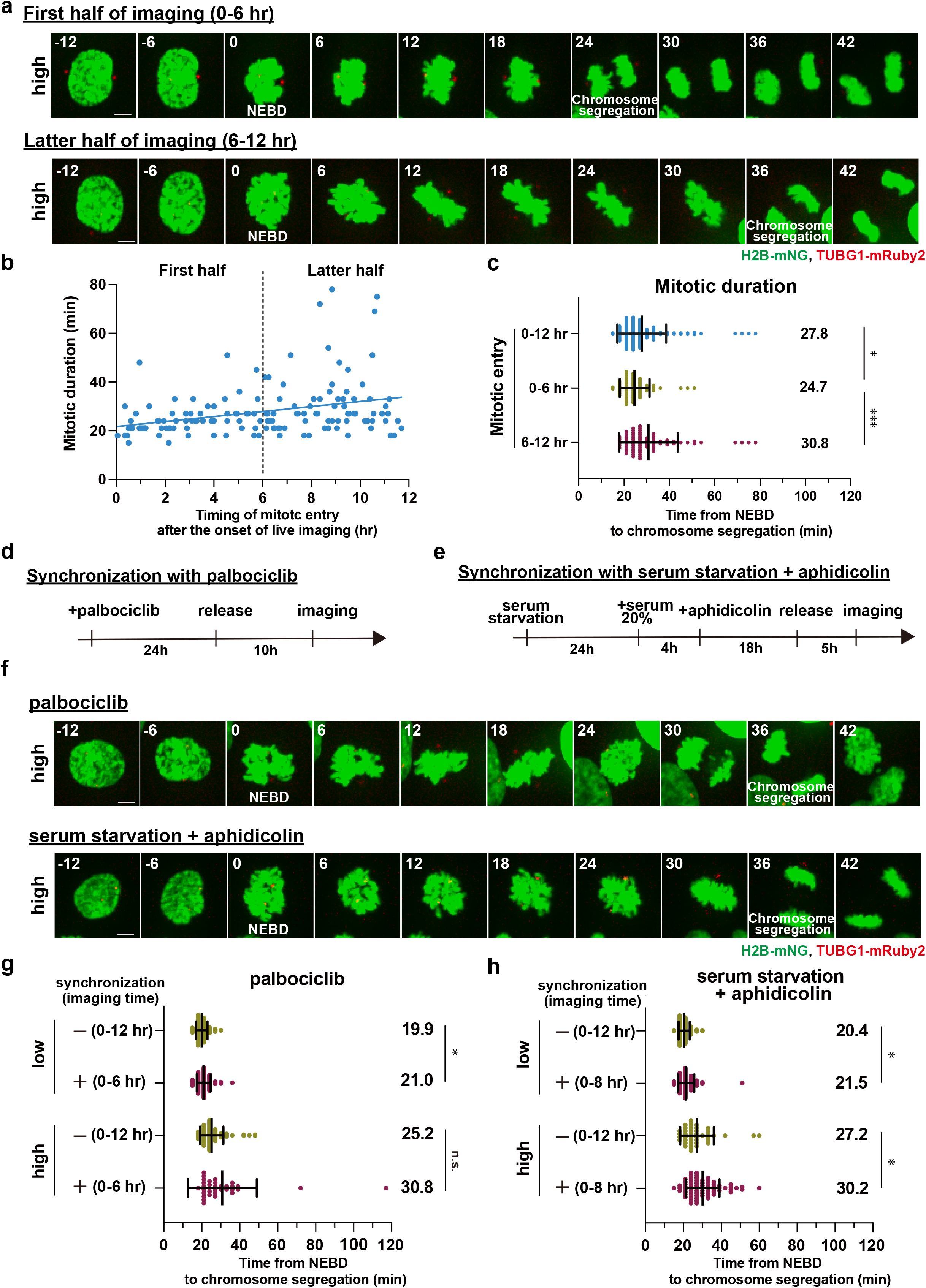
Reducing the total acquisition time by cell-cycle synchronization does not alleviate the light-induced mitotic prolongation. **a**, Time-lapse imaging of mitotic cells under the high condition (3-min intervals). Representative still images in the first and latter halves of the total acquisition time are shown. T=0 is designated as NEBD (time shown in min). Scale bar, 5 μm. **b**, The correlation between the time point of mitotic entry after the start of live imaging (x-axis, hours) and mitotic duration (y-axis, minutes) from **a**. NEBD was defined as the timing of mitotic entry. The regression line is shown. The vertical dotted line indicates the boundary of the first and latter 6 hr-halves after the onset of live imaging. n=150 cells from six independent experiments. **c**, Quantification of mitotic duration from **a**. **d**, Experimental procedure for cell-cycle synchronization with palbociclib treatment. **e**, Experimental procedure for cellcycle synchronization with serum starvation plus aphidicolin treatment. **f**, Timelapse imaging of mitotic cells synchronized with 300 nM palbociclib or with serum starvation plus 1 μM aphidicolin under the high illumination conditions (3-min intervals). Representative still images are shown. T=0 is designated as NEBD (time shown in min). Scale bar, 5 μm. **g**, Quantification of mitotic duration in the indicated conditions for the synchronization with palbociclib from **f**. In the control and synchronized cells, live imaging was performed for 12 hr and 6 hr, respectively. n>30 cells from three independent experiments. **h**, Quantification of mitotic duration in the indicated conditions for the synchronization with serum starvation plus aphidicolin treatment from **f**. In the control and synchronized cells, live imaging was performed for 12 hr and 8 hr, respectively. n>40 cells from three independent experiments. In **c**, **g**, and **h**, data are mean ± S.D., and *P* values were calculated by Mann-Whitney *U*-test. *P < 0.05, **P < 0.01, ***P < 0.001. n.s.: not significant.

Whereas shortening the total acquisition time of live imaging is one approach to reduce the pre-mitotic light exposure time of cells, this limits the number of mitotic cells that could be recorded in a single experiment. In order to increase the number of the cells that enter mitosis within a narrow time window, we examined two different cell-cycle synchronization methods to enrich premitotic cells prior to live cell imaging (Fig. 2d, e). The cell cycle progression of RPE1 cells has been shown to be highly synchronized after release from G1 arrest induced with palbociclib, a CDK4/6 inhibitor (Trotter and Hagan, 2020). Since we observed that the majority of the cells entered mitosis at 13-14 hr after the G1 release (Fig. S1), live cell imaging was started 10 hr after the palbociclib washout. Unexpectedly, even though the cell-cycle synchronization allowed us to shorten the total acquisition time (6 hr), compared to the control unsynchronous culture (12 hr), this approach did not suppress the light-induced mitotic prolongation (Fig. 2f, g). Furthermore, even under the low-light illumination, the synchronized cells showed a slight prolongation of mitosis. This result suggests that cell-cycle synchronization with palbociclib itself is potentially harmful to the cells.

We therefore explored another cell-cycle synchronization method, in which RPE1 cells were first arrested in G0 by serum starvation, released, then again arrested in G1/S with a low concentration of aphidicolin, and finally released to undergo mitosis in an accurately synchronized manner (Shi et al., 2007; Gheghiani et al., 2017). Live cell imaging was started 5 hr after the washout of aphidicolin to efficiently capture mitotic cells within a limited time window (Fig. 2e). Quantification analysis revealed that this synchronization approach also does not alleviate and even enhances the light-induced mitotic prolongation (Fig. 2f, h). Similar to the case of palbociclib treatment, cell-cycle synchronization with this two-step method resulted in a slight prolongation of mitosis even under the low-light illumination. Taken together, these data indicate that reducing the pre-mitotic light exposure time by cell-cycle synchronization does not alleviate the mitotic phototoxicity in fluorescence imaging with RPE1 cells, likely due to other toxic side-effects contributed by the chemical treatments.

### An antioxidant screen identifies ascorbic acid as a potent agent to prevent phototoxicity in mitosis

In order to reduce the phototoxicity in mitosis during fluorescence live cell imaging, we next examined the approach of adding antioxidants to the imaging medium. To identify the antioxidants that specifically protect mitotic cells from photodamage by high-light illumination, we performed a small screen of antioxidative agents known to reduce cellular phototoxicity (Bogdanov et al., 2012; Douthwright and Sluder, 2017; Natoli et al., 2016; Wrona et al., 2004; Wäldchen et al., 2015). Each antioxidant was examined in two different concentrations, the lower of which was chosen based on what is commonly used in live cell imaging experiments. Among the six compounds tested, we found that neither Trolox, nor zeaxanthin, sodium pyruvate, α-tocopherol or Rutin is able to prevent the light-induced mitotic prolongation (Fig. 3a, b, c, d, e, and f). This is surprising since Trolox has been shown previously to suppress the delay of mitotic entry caused by high-light illumination in RPE1 cells (Douthwright and Sluder, 2017) (Fig. 3a, b). In contrast, the addition of ascorbic acid to the imaging medium at 500 μM almost completely restored the duration of mitosis in the highlight condition to the values observed under low-light illumination (from 28.8 min to 19.8 min on average, Fig. 3a, g). Additionally, the mitotic duration was shorter in the presence of 500 μM ascorbic acid than that of 250 μM, suggesting a dose-dependent photoprotective effect of this antioxidant. Moreover, we observed that the treatment with 500 μM ascorbic acid significantly suppressed the delays in chromosome congression and centrosome separation shown to be induced by high amounts of light exposure (Fig. 3h, i). These results revealed that ascorbic acid, but not other commonly used antioxidants, is able to govern protection from phototoxicity specifically to mitotic cells.

**Fig.3:**
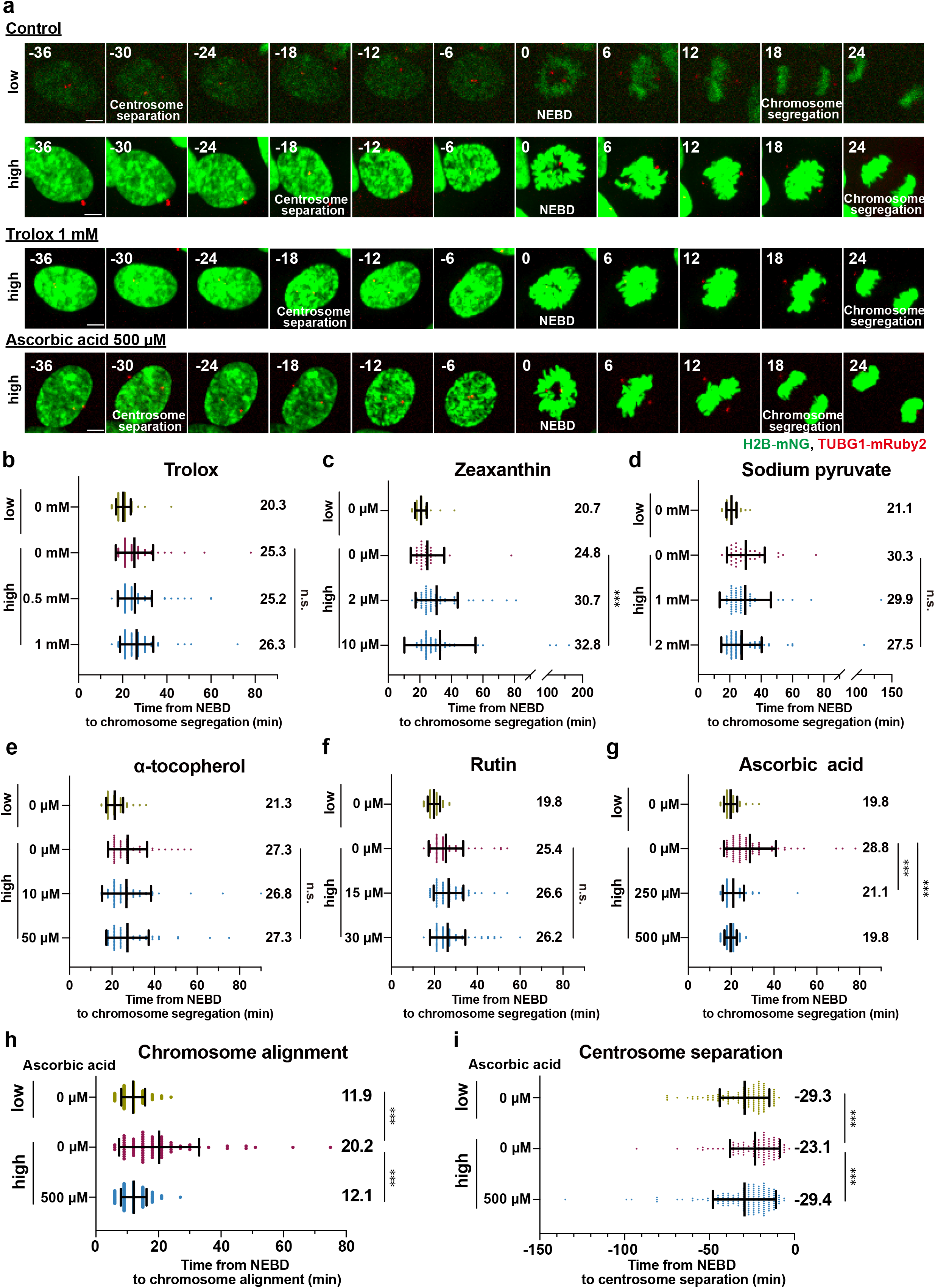
An antioxidant screen identifies ascorbic acid for reducing phototoxicity to mitotic cells. **a**, Time-lapse imaging of mitotic cells in the presence of antioxidants. Representative still images of control cells or cells imaged in the presence of 1 mM Trolox, or 500 μM ascorbic acid, are shown (3-min intervals). Images with different settings of brightness and contrast are shown. T=0 is designated as NEBD (time shown in min). Scale bar, 5 μm. **b-g**, Quantification of mitotic durations in the presence of the following antioxidants from **a**: **b**: Trolox, **c**: Zeaxanthin, **d**: Sodium pyruvate, **e**: α-tocopherol, **f**: Rutin, and **g**: Ascorbic acid. n>30 cells from three independent experiments. **h**, Quantification of the time required for chromosome alignment in the absence and the presence of 500 μM ascorbic acid from **a**. The time from NEBD to chromosome alignment at the metaphase plate was measured. n>70 cells from three independent experiments. **i**, Quantification of the timing of centrosome separation in the absence and the presence of 500 μM ascorbic acid from **a**. The time from centrosome separation (>4 μm inter-centrosomal distance) to NEBD was measured. n>90 cells from five independent experiments. In **b**-**i**, data are mean ± S.D., and *P* values were calculated by Mann-Whitney *U*-test. ***P < 0.001, n.s.: not significant.

### Ascorbic acid is not cytotoxic at the concentration required for prevention of mitotic phototoxicity

We next examined whether the concentration of ascorbic acid exerting the photoprotective effect could be generally harmful to the cells. To this end, RPE1 cells were treated with various concentrations of ascorbic acid for 72 hours and cell viability was subsequently determined by MTT assay for each condition (Fig. 4a, b). A dose-dependent curve of ascorbic acid on cell viability was plotted and the 50% inhibitory concentration value (IC50) was calculated as 3.38 mM. Whereas ascorbic acid concentrations above 1 mM reduced the viability of RPE1 cells, lower concentrations had no effect. Fluorescence live cell imaging in the low-light illumination regime confirmed that the accuracy of chromosome segregation was not affected in the presence of ascorbic acid at 500 μM (Fig. 4c, d). Thus, these data indicate that the phototoxicity to mitosis can be safely suppressed by the use of ascorbic acid at a non-cytotoxic dose.

**Fig.4:**
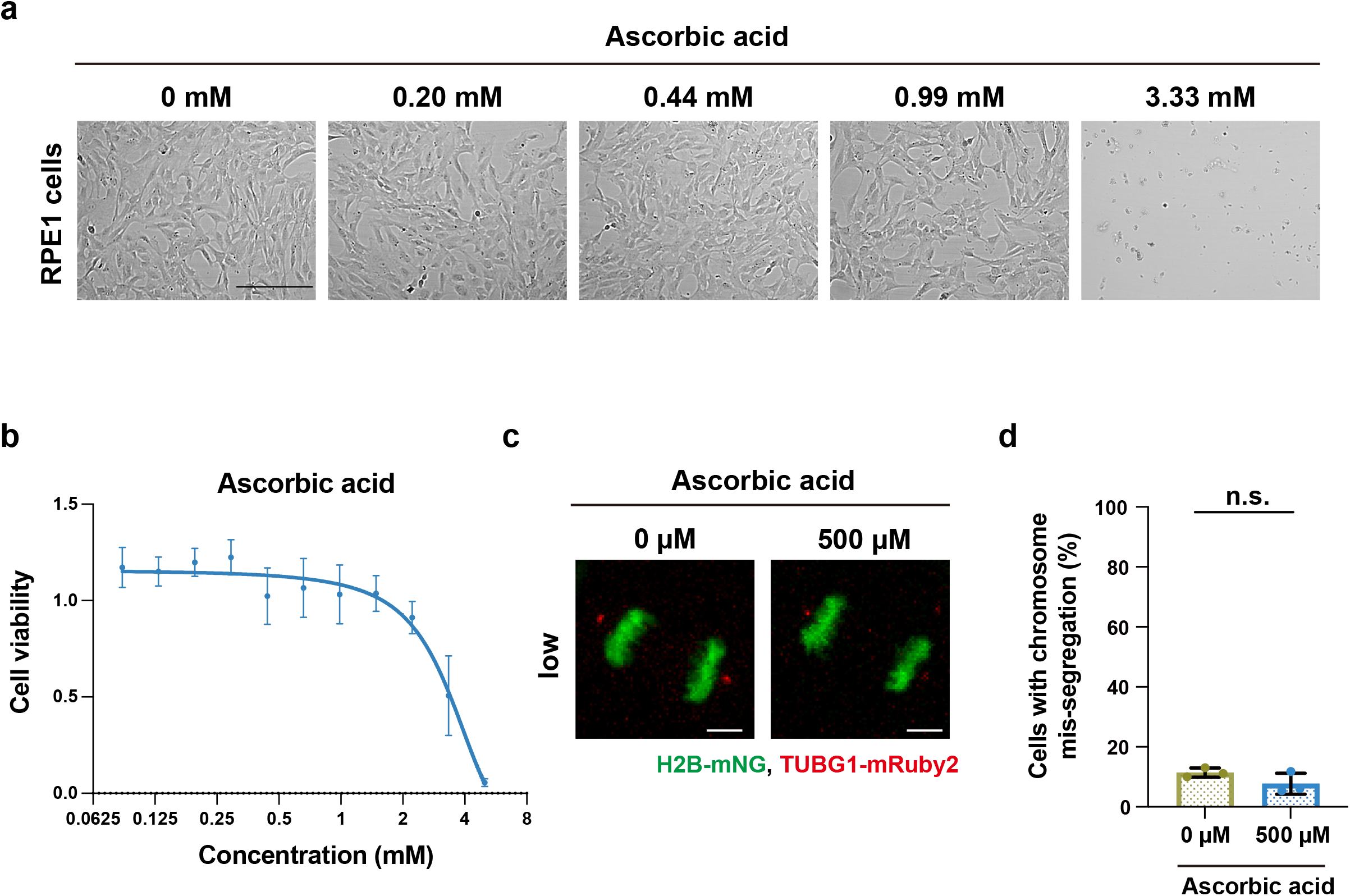
Ascorbic acid at concentrations adequate to prevent the mitotic phototoxicity is not cytotoxic. **a**, Cell viability assay of RPE1 cells in the presence of various concentrations of ascorbic acid. Representative bright-field images for the indicated conditions are shown. Scale bar, 500 μm. **b**, Quantification of dose-dependent effect of ascorbic acid on cell viability assessed by MTT assay. Cell viability was normalized with 0 mM ascorbic acid. n=5 biological replicates for each concentration. Values are given as mean ± S.D. and the interpolation curve is shown. **c**, Chromosome segregation of RPE1 cells stably expressing H2B-mNG and TUBG1-mRuby2 at the indicated concentrations of ascorbic acid in the low-light condition. Representative images of anaphase cells are shown. Scale bar, 5 μm. **d**, Quantification of the percentage of chromosome mis-segregation from n>90 cells from three independent experiments. Data are mean ± S.D., and *P* values were calculated by two-tailed unpaired Student’s t-test. n.s.: not significant.

### The presence of ascorbic acid allows very high temporal resolution imaging of mitosis without obvious photodamage

High temporal resolution imaging is necessary to capture the rapid dynamics of mitotic processes. However, shortening the interval between two frames in time-lapse imaging results in more intensive light exposure of the cell sample, which in turn leads to higher phototoxicity. We therefore hypothesized that adding ascorbic acid to the imaging media could alleviate the severe phototoxicity in mitosis caused by the shortening of imaging intervals. To test this idea, we set a new imaging condition with 30-sec intervals, which is equivalent in light intensity to the low condition with 3-min intervals (Fig. 5a). Whereas the short-interval condition allowed us to capture the fine movement of chromosomes in mitosis, it resulted in a significantly longer duration of mitosis compared to the condition with 3-min imaging intervals (Fig. 5b, c). In addition, the cells that entered mitosis at later time points during the imaging showed longer mitotic durations in the short-interval condition, as seen in the high-light illumination condition with 3-min intervals (Fig. 5d and 2b). Remarkably, the addition of ascorbic acid to the imaging media restored the precise duration of mitosis (from 28.0 min to 21.2 min on average) even in the short-interval condition (Fig. 5b, c). This outstanding photoprotective property of ascorbic acid was observed throughout the entire time-lapse acquisition (~12 hr), abolishing the effect of increasing mitotic duration with accumulating amount of blue-light exposure (Fig. 5d). Thus, these data indicate that the use of ascorbic acid enables to capture the accurate mitotic processes at very high temporal resolution of fluorescence live imaging.

**Fig.5:**
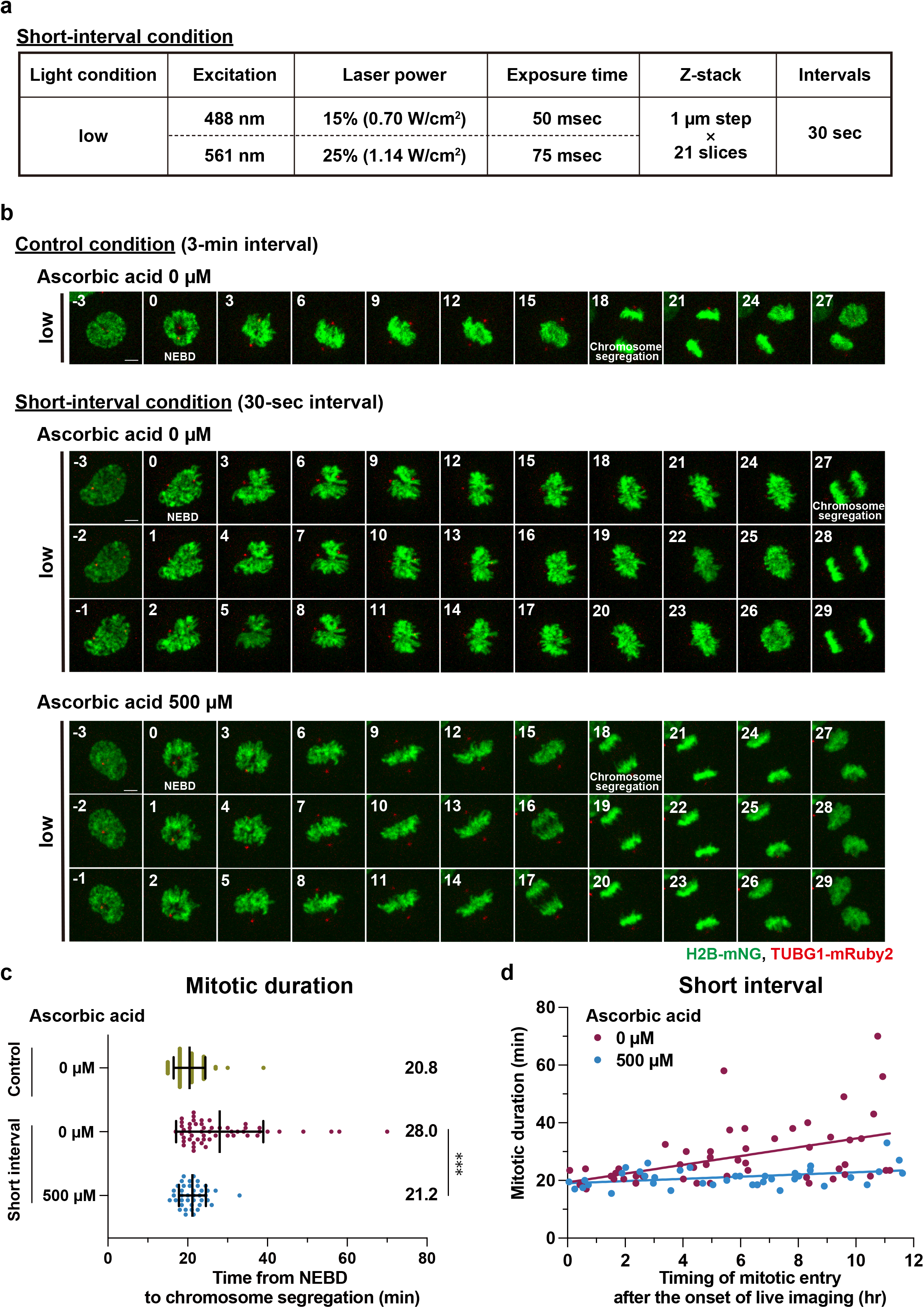
The addition of ascorbic acid to imaging buffer enables very high temporal resolution imaging of mitosis without obvious photodamage. **a**, The imaging condition used for live cell imaging with 30-sec intervals. **b**, Timelapse imaging of mitotic cells in the indicated conditions. Representative still images with different settings of brightness and contrast are shown. T=0 is designated as NEBD (time shown in min). Scale bar, 5 μm. **c**, Quantification of mitotic duration from **b**. The time from NEBD to chromosome segregation was measured. n≥40 cells from four independent experiments. Data are mean ± S.D., and *P* values were calculated by Mann-Whitney *U*-test. ***P < 0.001. **d**, The correlation between the timing of mitotic entry (NEBD) after the start of live imaging (x-axis) and mitotic duration (y-axis) in the short-interval condition from **b**. The regression lines for the indicated conditions are shown.

## Discussion

In this study, we demonstrate that a submillimolar concentration of ascorbic acid can be safely applied to reduce phototoxicity-induced mitotic defects during fluorescence live-cell imaging. We first analyzed the phototoxicity of excitation light on mitotic dynamics and observed that, consistent with a previous report, the duration of mitosis was significantly prolonged by high dose illumination with excitation light of 488 nm. In particular, the process of chromosome alignment at the metaphase plate was delayed. Blue light illumination has been shown to induce ROS-mediated DNA damage, such as oxidation of DNA bases and DNA double-strand breaks (DSBs) (Kielbassa et al., 1997; Cheng et al., 2021). Moreover, DSBs generated during mitosis are reported to cause the delay of chromosome alignment and the activation of spindle assembly checkpoint, resulting in prolongation of the total mitotic duration (Choi et al., 2008; Mikhailov et al., 2002; Bakhoum et al., 2017). Similarly, the light-induced prolonged mitosis observed under our experimental conditions may be attributed to DSBs induced by high-light illumination. Besides mitotic duration and chromosome alignment, our analysis additionally revealed that the timing of centrosome separation is another photo-sensitive process in mitosis. Centrosome separation is regulated by a mitotic kinase cascade consisting of PLK1, NEK2, and others (Mardin et al., 2012). Previous reports have shown that DNA damage inactivates these mitotic kinases leading to the inhibition of centrosome separation (Fletcher et al., 2004; Zhang et al., 2005). Thus, the light-induced DNA damage is likely to be a common cause of the here observed mitotic defects in fluorescence live cell imaging.

To alleviate the phototoxicity to mitosis in fluorescence live cell imaging, we examined the effect of reduced light exposure prior to entering mitosis by introducing a cell-cycle synchronization step before the start of acquisition experiment. Although two different synchronization methods allowed shortening of the total light exposure time before mitosis, these approaches did not suppress the light-induced mitotic prolongation, likely due to cytotoxic side-effects. Aphidicolin, used in the two-step synchronization method, is known to cause replication stress even at low concentrations (Lukas et al., 2011; Koundrioukoff et al., 2013). Cells with mild replication stress can escape from the S phase checkpoint and progress into mitosis, although they tend to exhibit prolonged mitosis (Wilhelm et al., 2014). Therefore, the cytotoxic effect of aphidicolin may amplify the phototoxic effect instead of relieving it. Even in the case of synchronization with palbociclib treatment, replication stress is a non-negligible factor. Long-term treatment of palbociclib has been shown to induce replication stress by downregulating the components of the replisome (Crozier et al., 2022). While 24 hours of palbociclib treatment is not likely sufficient to induce obvious replication stress, it is possible that mild replication stress by palbociclib treatment, together with high-light illumination, synergistically induces mitotic prolongation. Thus, our work suggests that the approaches of cell-cycle synchronization are not suitable for alleviating the light-induced mitotic prolongation in fluorescence live cell imaging due to their known activation of replication stress.

Our antioxidant screen identified ascorbic acid as a potent supplement that can be added to the imaging media to suppress the light-induced mitotic prolongation. In contrast, other antioxidants known to prevent ROS activity and cellular phototoxicity did not alleviate the mitotic phototoxicity in this study. This is surprising because some of these antioxidants have been shown to inhibit ROS-mediated DNA DSBs (Safaei et al., 2018), suggesting that the outstanding effect of ascorbic acid may not solely rely on the ROS scavenging activity. Interestingly, it has been shown that ascorbic acid possesses the additional property of compacting the higher-order structure of DNA through direct interaction with DNA, which suppresses photo-induced DSB formation (Yoshikawa et al., 2003; Yoshikawa et al., 2006). Therefore, this unique property of ascorbic acid, together with the ROS scavenging activity, may be responsible for its outstanding effect in alleviating the light-induced mitotic prolongation. Taken together, this study revealed ascorbic acid as the optimal antioxidant to reduce the mitotic phototoxicity occurring during prolonger light exposure, and thus offers a practical solution for low-phototoxicity fluorescence live cell imaging with very high temporal resolution.

## Acknowledgments

We thank Mariya Genova for the proofreading of the manuscript and the Kitagawa lab members for technical supports and helpful discussions. This work was supported by JSPS KAKENHI grants (Grant numbers: 18K06246, 19H05651, 20K15987, 20K22701, 21H02623, 22H02629) from the Ministry of Education, Science, Sports and Culture of Japan, the PRESTO program (JPMJPR21EC) of the Japan Science and Technology Agency, Takeda Science Foundation, The Uehara Memorial Foundation, The Research Foundation for Pharmaceutical Sciences, Koyanagi Zaidan, The Kanae Foundation for the Promotion of Medical Science, Kato Memorial Bioscience Foundation, and Tokyo Foundation for Pharmaceutical Sciences.

## Author contributions

S.H. conceived and designed the study. T.H. performed all the experiments. M.F. and T.C. provided suggestions. T.H., S.H., and D.K. analyzed the data. T.H., S.H., and D.K. wrote the manuscript. All authors contributed to discussions and manuscript preparation.

## Material and methods

### Cell culture and cell-cycle synchronization

RPE1 cells were grown in Dulbecco’s Modified Eagle’s Medium F-12 (DMEM/F-12) with 10% FBS and 1% penicillin/streptomycin, and cultured at 37°C in a humidified 5% CO2 incubator. For cell-cycle synchronization with palbociclib, cells were treated with palbociclib (300nM) for 24 hr and released by replacing with fresh media. For the starvation and aphidicolin synchronization, cells were first serum starved with medium containing 0.1% FBS for 24 hr and subsequently released by replacing with medium containing 20% FBS. After 4 hr of the release, cells were treated with aphidicolin (1 μM) for 18 hr and released by replacing with fresh media.

### Antioxidants

The following antioxidants were used in this study: Trolox (Vector Laboratories), Zeaxanthin (Santa Cruz), α-tocopherol (Nacalai Tesque), L-Sodium Pyruvate (Nacalai Tesque), Rutin (Nacalai Tesque), and Ascorbic acid (Nacalai Tesque). Ascorbic acid (100 mM) was prepared in water and stored at −20°C. Zeaxanthin (2 mM), α-tocopherol (10 mM), and Rutin (10 mM) were prepared in DMSO and stored at −20°C. Trolox (100 mM) and Sodium Pyruvate (100 mM) were stored at 4°C. Each antioxidant was diluted in the culture medium at a final concentration before live cell imaging.

### Fluorescence live cell imaging

For fluorescence live cell imaging, RPE1 cells stably expressing mNG-H2B and TUBG1-mRuby2 were cultured in 35-mm glass-bottom dishes (Greiner-bio-one, #627870) at 37°C in a 5% CO2 atmosphere. The growth medium was replaced with modified medium containing each antioxidant 3 hr before live cell imaging. A spinning disk confocal scanner box, the Cell Voyager CV1000 (Yokogawa Electric Corp) equipped with a 40× oil-immersion objective and a back-illuminated EMCCD camera was used for live cell imaging. Images were acquired as z-stacks with a 40× 1.30 NA oil-immersion lens every 3 min or 30 sec for 12 hr. Maximum intensity projections of representative images were created using Fiji. The power of excitation light was measured using laser power meter 3664 (HIOKI).

### MTT assay

RPE1 cells were seeded at a density of 1×10^4^ cells/ml in a 96-well plate. After 24 hr, the medium was replaced with fresh one containing each concentration of ascorbic acid. The plate was then incubated for 72 hr, and the medium was replaced with medium containing MTT (ThermoFisher, 0.5 mg/ml final concentration). After culture for 3 hr, the medium was discarded and MTT formazan crystals were dissolved by adding isopropanol. The optical density was read at 600 nm. Cell viability was calculated by the normalization of optical density to the control treatment. Representative images were acquired using CQ1 Benchtop High-Content Analysis System equipped with a 10× objective and a sCMOS camera (Yokogawa Electric Corp).

### Statistics analysis

Statistical comparison between the data from different groups was performed in PRISM v.9 software (GraphPad) using Mann-Whitney *U*-test or two-tailed unpaired Student’s t-test. P values <0.05 were considered statistically significant. All data shown are mean ± S.D. The sample size is indicated in the figure legends.

